# Investigating the cytotoxic effects of the venom proteome of two species of the *Viperidae* family (*Cerastes cerastes* and *Cryptelytrops purpureomaculatus*) from various habitats

**DOI:** 10.1101/449728

**Authors:** Cenk Serhan Ozverel, Maik Damm, Benjamin-Florian Hempel, Bayram Göçmen, Robert Sroka, Roderich D. Süssmuth, Ayse Nalbantsoy

## Abstract

Animal secretions are of great interest in terms of drug development due to their complex protein and peptide composition. Especially, in the field of therapeutic medications such as anti-cancer drugs snake venoms receive attention. In this study we report of two *Viperidae* species from various habitats with a particular focus on the cytotoxic potential along with the decomplexation of the venom proteome: the horned desert viper (*Cerastes cerastes*), native to desert regions of North Africa and the mangrove pit viper (Cryptelytrops purpureomaculatus), found in coastal forests of Southeast Asia. Initial cytotoxic screenings of the crude venoms revealed diverse activity, with the highest effect against SHSY5Y human glioblastoma carcinoma cells compared to other cancerous and non-cancerous cell lines. In-depth cytotoxicity studies of SHSY5Y cells with purified venom fractions revealed dimeric disintegrins from *C. cerastes* venom which exerted a high cytotoxic activity with IC_50_ values from 0.11 to 0.58µM and the disintegrins-like effect on SHSY5Y morphology was observed due to cell detachment. Furthermore, two polyproline BPP-related peptides, one PLA_2_ and a peptide-rich fraction were determined for *C. purpureomaculatus* with moderate IC_50_ values between 3-51µM. Additionally, the decryption of the venom proteomes by snake venomic mass spectrometry and comparison of same species from different habitats revealed slight differences in the composition.

## 1 INTRODUCTION

Secretions from animals have been versatile studied in recent decades for treating several diseases, like Leishmania, hypertension, Alzheimer’s diseases, congestive heart failure, as well as different types of cancer (Chan et al., 2016; Macedo et al., 2015; Smith and Vane, 2003; Vyas et al., 2013). Preliminary studies suggest especially for snake venoms great potential of anti-cancer properties (Das et al., 2013; Jain and Kumar, 2012; Konshina et al., 2011; Li et al., 2018; Marsh and Williams, 2005). One of the interesting aspects of snake venoms in cancer treatment studies is the well-known ability of isolated toxin components to interact with membranes and thus to inhibit the general cell migration and proliferation (Kitchens and Eskin, 2008; Vyas et al., 2013). Snake venoms as valuable secretions are composed in general of manifold proteins and peptides with remarkable biological activities (Aird, 2005; Devi, 2013; Fry, 2015; Mackessy, 2010; Vyas et al., 2013). They comprise proteolytic enzymes, non-cholinesterase enzymes, thrombin, thrombin-like enzymes with anticoagulant activities, collagenases, phospholipases, and others (Antunes et al., 2010; Biardi et al., 2006; Kitchens and Eskin, 2008; Sales and Santoro, 2008; Vieira Santos et al., 2008). These venom components with enzymatic activities are vital for snake survival tactics, including defense, immobilization and digestion of preys (Casewell et al., 2013; Das et al., 2013). Especially, the multifaceted venom proteomes of vipers, with their high cytotoxic and coagulopathic potential are considered to bear great potential for development of medical treatments (Fry, 2015; Harvey, 2014). *In vitro* studies on crude venoms and purified snake proteins showed cytotoxic effects against various cancerous human cell lines (Bradshaw et al., 2016; Kakanj et al., 2015; Nalbantsoy et al., 2017). The venom content was shown to exert their activities e.g. via blockage of integrins, induction of apoptosis or necrosis pathways and disruption of the cell cycle (Calderon et al., 2014; Göçmen et al., 2015a; Sciani and Pimenta, 2017; Yalcın et al., 2014; Zainal Abidin et al., 2018). This correlates to some extent with chemotherapeutic agents that are currently being used in clinics, making venoms attractive sources of future anti-cancer agents (Calderon et al., 2014; Meyer et al., 2004; Nalbantsoy et al., 2017; Suzergoz et al., 2016).

The occurrence of the same snake species in different geographical locations could have effects on the venom content, i.a. based on divergence of the prey (Bazaa et al., 2005; Tashima et al., 2008). It is known that, venoms from the same species may contain differences in their composition due to local regions as well as following narrow diets on the individual and population level (Barlow et al., 2009; Chijiwa et al., 2000; Daltry et al., 1996; Gibbs and Mackessy, 2009; Greene, 1997). Additionally, variations in the ecological pressures, abundancy, availability and types of prey may affect the body size and body mass of snakes (Barlow et al., 2009; Schwaner and Sarre, 1988). Representatives of the *Viperidae* family can be found in various geographical locations with highly diverse climatic conditions around the world, for instance the European/Anatolian *Vipera ammodytes*, the American *Crotalus durissus*, the North African *Cerastes cerastes* (*C*. cerastes) as well as the Asian *Cryptelytrops purpureomaculatus* (*C. purpureomaculatus*) (Fahmi et al., 2012; Göçmen et al., 2015b; Tomović, 2006; Vanzolini and Calleffo, 2002; Vogel et al., 2012). Therefore, the broad distribution of *Viperidae* within different environmental conditions makes them a good study object for analyzing venom compositions and assess variations of their content (Chippaux et al., 1991).

The variations of cytotoxic constituents between *Viperidae* species venoms is directly linked to the proteomic content, which in turn depends on the habitat and above mentioned factors (Barlow et al., 2009; Greene, 1997; Schwaner and Sarre, 1988; Tashima et al., 2008). In this study, we focused on the cytotoxic properties, we quantified the composition and identification of these cytotoxins of two viper species located in different habitats: the Egypt *C. cerastes* and the Thai *C. purpureomaculatus* (see **Figure 1**). The horned desert viper *C. cerastes* is mainly hunting during the night and specialized at rodents, lizards and other small animals that are affected by its venom composition (Al-Sadoon and Paray, 2016; Calvete et al., 2002; Dekhil et al., 2003). Previous studies on the venom proteome of *C. cerastes* showed the presence of disintegrins (DI), procoagulant serine proteinase cerastocytin, phospholipase A_2_ (PLA_2_), C-type lectin-like proteins (CTL) and metalloproteinases (svMP) (Bazaa et al., 2005; Calvete et al., 2002; Dekhil et al., 2003; Fahmi et al., 2012). In contrast, the *C. purpureomaculatus* is mainly found in more tropical regions of Southeast Asia, whereby its prey is composed of warm blooded species and is less species-divergent compared to *C. cerastes* (Gopalakrishnakone et al., 2015; Leong et al., 2014). The venom of *C. purpureomaculatus* contains svMP, PLA2, serine proteases (svSP), phosphodiesterases (PDE), 5′-nucleotidases, and phospholipase B, while a bite is not fatal to humans but leads to pain and swelling (Fahmi et al., 2012; Mong and Tan, 2016; Zainal Abidin et al., 2016). Previous bioactivity studies on the venom of *C. purpureomaculatus* showed thrombin-like, hemorrhagic, anti-coagulant activities, with higher toxicity relative to other Asian viper species (Tan et al., 1989).

**Figure 1:**
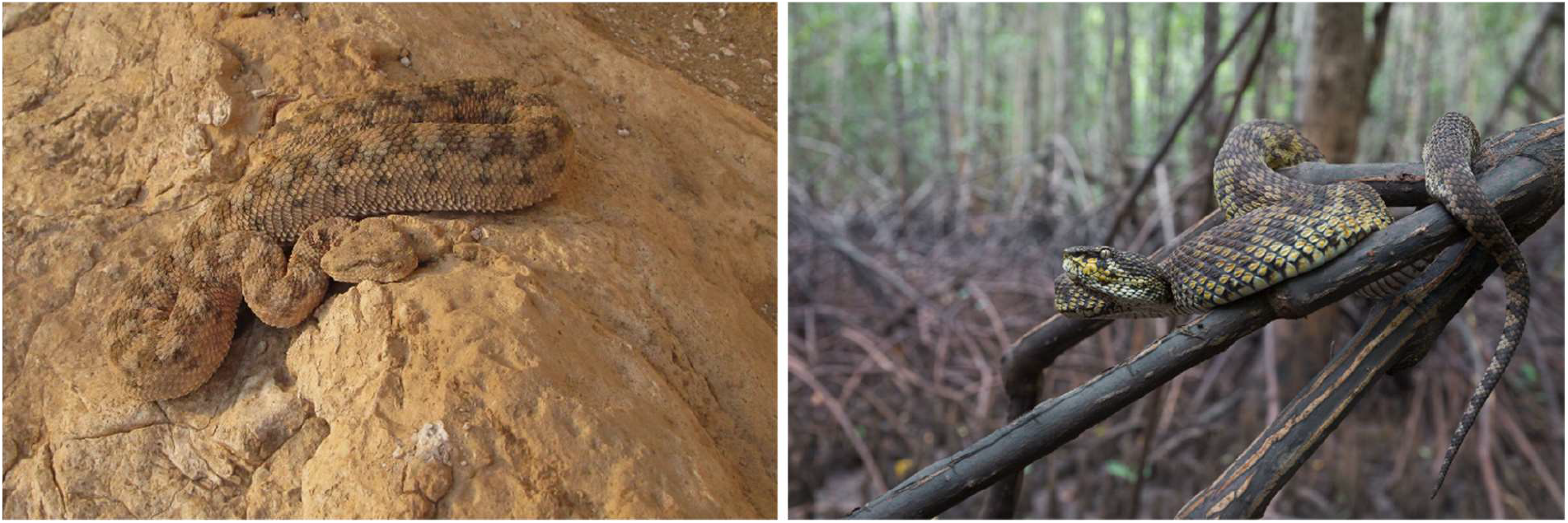
Exemplary representation of *Cerastes cerastes* and *Cryptelytrops purpureomaculatus* in their natural habitats. *C. cerastes* (left) is shown in the desert and rocky regions of Egypt and *C. purpureomaculatus* in the mangrove forests of Thailand.

In this study, crude venom samples from these two viper species were tested on a panel of eight cancerous cell lines together with one non-cancerous cell line. Further, we identified venom fractions via bottom-up and intact mass profiling in order to investigate the potentially active peptides as well as to determine the venomic composition.

## 2 MATERIALS AND METHODS

### 2.1 Source of animals and venom collection

Individuals of *C. cerastes* (horned desert viper) from Egypt and C. *purpureomaculatus* (mangrove pit viper) from Thailand were collected in their naturally-occurring habitats and kept in Turkey illegally. Both vipers are retained by the Republic of Turkey, Ministry of Forestry and Water Affairs upon a notification and given to Prof. Dr. Bayram Göçmen with a special permission for studies. The study was approved by the Ege University, Local Ethical Committee of Animal Experiment (Approval Number: 2017/020).

Crude venoms from both species were extracted by using a paraffin-covered laboratory beaker without applying any pressure on the venom glands. Venoms collected from both species were centrifuged at 2000 xg for 10 min at 4 °C for the elimination of debris. Supernatants were collected after centrifugation and venom samples were directly frozen at -80 °C, lyophilized and stored at 4 °C.

### 2.2 Determination of protein content

Protein concentrations were determined from diluted venom samples (2mg/mL) in ultra-pure water by Bradford assay (Bradford, 1976) using a UV/Vis spectrophotometer (Thermo-Scientific, Darmstadt, Germany) at a wavelength of λ =595 nm. Bovine serum albumin was used as a reference.

### 2.3 Cell culture and *in vitro* cytotoxicity assay

Eight human cancerous and one non-cancerous cell lines were used for the determination of cytotoxicity. The tested cell lines were as followed: SHSY5Y (neuroblastoma cell line), MCF-7 (breast adenocarcinoma), HeLa (cervix adenocarcinoma), A-549 (alveolar adenocarcinoma), CaCo-2 (colon colorectal adenocarcinoma), MDA-MB-231 (breast adenocarcinoma), 253-JBV (bladder carcinoma), U87MG (glioblastoma astrocytoma) as cancerous cell lines and HEK 293 (embryonic kidney cells) were used as non-cancerous cell line. Cell lines were purchased from ATCC (Manassas, VA, USA). All cell lines were cultivated in Dulbecco’s modified Eagle’s medium F12 (DMEM/F12), supplemented with 10% fetal bovine serum (FBS), 2mM/L glutamine, 100U/mL of penicillin and 100mg/mL of streptomycin (Gibco, Visp, Switzerland).

The cells were incubated at 37 °C in a humidified atmosphere of 5% CO_2_. The cytotoxicity of crude venoms and fractions were determined by using a modified MTT [3-(4,5-dimethyl-2-thiazolyl)-2,5-diphenyl-2H-tetrazoliumbromide)] assay (Mosmann, 1983; Ozen et al., 2015), which detects the activity of the mitochondrial reductase of viable cells. The assay principle is based on the cleavage of MTT that forms formazan crystals by cellular succinate-dehydrogenases in viable cells. Insoluble formazan crystals were dissolved by the addition of DMSO. In order to perform the cytotoxicity assay, all cell lines were cultivated for 24 h in 96-well microtiter plates with an initial concentration of 1×10^5^ cells/mL. Afterwards, the cultured cells were treated with different concentrations of crude venom or fractions and incubated for 48 h at 37 °C. The optical density (OD) was measured in triplicates at λ=570 nm (with a reference wavelength λ=690 nm) by UV/Vis spectrophotometry (Thermo, Bremen, Germany). Percentages of surviving cells in each culture were determined after incubation with venom. The viability (%) was determined by the following formula with an absorbance A:

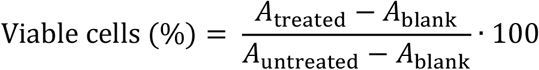

### 2.4 Determination of half maximal inhibitory concentration (IC50)

The half maximal inhibition of growth (IC_50_) values were calculated by fitting the data to a sigmoidal curve and using a four-parameter logistic model and presented as an average of three independent measurements. The IC_50_ values were reported at 95% confidence interval. Calculations were performed via Prism 5 software (GraphPad5, San Diego, CA, USA). The values of the blank wells were subtracted from each well of treated and control cells and IC_50_ were calculated in comparison to untreated controls.

### 2.5 Morphological studies

The morphological changes of the cells were observed via an inverted microscope (Olympus, Tokyo, Japan). All groups treated with different dosages were compared to the control group following 48 h treatment.

### 2.6 Statistics

All data were studied as triplicates and are presented as mean ± standard error of mean (SEM). For the calculation of IC_50_ values Prism 5 software (GraphPad5, San Diego, CA, USA) was used to analyze variance (standard deviation calculation).

### 2.7 Intact mass profiling (IMP)

Crude venom was dissolved to a final concentration of 10mg/mL in 5 µL aqueous formic acid (HFo, 1% v/v) and centrifuged at 20,000 xg for 5 min to sediment the cell debris. The supernatant was diluted with 25 µL citrate buffer (0.1 M, pH 4.3) and submitted to an intact mass profiling: HPLC-high-resolution (HR) ESI-MS/MS measurements were performed on a LTQ Orbitrap XL mass spectrometer (Thermo, Bremen, Germany) coupled to an Agilent 1260 HPLC system (Agilent, Waldbronn, Germany) using a Supelco Discovery 300 Å C18 (2 x 150 mm, 3 μm particle size) column. The elution was performed by a gradient of ultra-pure water with 0.1% HFo (v/v; buffer A) and ACN with 0.1% HFo (v/v; buffer B) at a flow rate of 1 mL/min. An isocratic equilibration (5% B) for 5 min was followed by a linear gradient of 5-40% B for 95 min, 40-70% B for 20 min, 70% B for 10 min and a re-equilibration with 5% B for 10 min.

ESI settings were: 11 L/min sheath gas, 35 L/min auxiliary gas, spray voltage 4.8 kV, capillary voltage 63 V, tube lens voltage 135 V, and capillary temperature 330 °C. The data-dependent acquisition (DDA) mode was used for MS/MS experiments with 1 μ scans and 1000 ms maximal fill time. The precursor ions were selected with a range of ±2 *m/z* and after two repeats within 10 s excluded with ±3 *m/z* for duration of 20 s. Three scan events were performed with a normalized CID energy of 30% and a HCD with 35% collision energy.

For the IMP, the MS1 spectra were inspected via the Xcalibur Qual Browser (Thermo Xcalibur 2.2 SP1.48) and the peak assignment was performed regarding the BU annotation, based on the use of analog columns. Deconvoluted isotopically resolved MS/MS spectra were generated by using the XTRACT algorithm of the Xcalibur Qual Browser. The protein assignment was done by comparison to the retention time of the HPLC runs. The mass comparison was performed manually.

### 2.8 Bottom-up (BU) venomics

Crude venom (4 mg, lyophilized) was dissolved to a final concentration of 20 mg/mL in aqueous 3% (v/v) ACN with 1% (v/v) HFo, centrifuged at 20,000 xg for 5 min and the supernatant was loaded onto a semi-preparative reversed-phase HPLC with a Supelco Discovery BIO wide Pore C18-3 column (4.6 x 150 mm, 3 μm particle size) using an Agilent 1260 Low Pressure Gradient System (Agilent, Waldbronn, Germany). Ultra-pure water with 0.1% (v/v) HFo (buffer A) and ACN with 0.1% (v/v) HFo (buffer B) was used with a flow rate of 1 mL/min. An isocratic equilibration (5% B) for 5 min was followed by the toxin elution with a linear gradient of 5-40% B for 95 min, 40-70% B for 20 min and 70% B for 10 min.

A DAD detector was used for absorbance measurements at λ=214 nm and fractions (1 mL) were automatically collected. Peaks of the chromatograms were manually pooled, vacuum dried (Thermo Speedvac, Bremen, Germany) and fractions were submitted to SDS-PAGE under reducing conditions (Laemmli, 1970). Coomassie-stained (Blue G250, Serva, Heidelberg, Germany) protein bands were cut, in-gel reduced with fresh dithiothreitol (100 mM DTT in 100 mM ammonium hydrogencarbonate, pH 8.3, for 30 min at 56 °C) and subsequent alkylated with fresh iodoacetamide (55 mM IAC in 100 mM ammonium hydrogencarbonate, pH 8.3, for 20 min at 25 °C). Proteins in the bands were in-gel digested with trypsin (Thermo, Rockfeld, IL, USA) with 6.7 ng/µL in 10 mM ammonium hydrogencarbonate with 10% (v/v) ACN, pH 8.3, for 18 h at 37 °C with 0.27 µg/band. Followed by peptide extraction with 100 µL aqueous 30% (v/v) ACN with 5% (v/v) HFo for 15 min at 37 °C. The supernatants were vacuum dried, re-dissolved in 20 µL aqueous 3% (v/v) ACN with 1% (v/v) HFo and submitted to LC-MS/MS analysis.

The analytics of tryptic peptides was performed using a reversed-phase Grace Vydac 218MSC18 column (2.1 x 150 mm, 5 μm particle size) under control of an Agilent 1260 HPLC system (Agilent Technologies, Waldbronn, Germany) with a flow rate of 0.3 mL/min. After an isocratic equilibration (5% B) for 1 min, the peptides were eluted with a linear gradient of 5-40% B for 10 min, 40-99% B for 3 min, washed with 99% B for 3 min and re-equilibrated in 5% B for 3 min. MS experiments were performed on an Orbitrap XL mass spectrometer (Thermo, Bremen, Germany) with R=15,000 at *m/z* 400 and maximum filling time of 200 ms for first product ion scans. MS/MS fragmentation of the most intense ion was performed in the LTQ using a collision-induced dissociation (30 ms activation time); the collision energy was set to 35%. The precursor ions were selected with a range of ±2 *m/z* and after two repeats within 10 s excluded with ±3 *m/z* for duration of 20 s.

LC-MS/MS data files (.raw) were converted to mascot generic format files (.mgf) via MSConvert GUI of the ProteoWizard package (http://proteowizard.sourceforge.net; version 3.0.10577) and annotated by DeNovo GUI 1.15.11 (Muth et al., 2014) with carbamidomethylated cysteine (+57.021464Da) as fixed modification. Variable modifications were acetylation of lysine (+42.010565Da) and phosphorylation of serine and threonine (+79.966331 Da).

The peptide sequences were matched against a non-redundant protein NCBI database of *Viperidae* (taxid: 8689) using BLASTP (http://blast.ncbi.nlm.nih.gov).

### 2.9 Relative toxin quantification

The percentage composition of the venom ingredients was calculated on a combination of the RP-HPLC chromatogram and SDS-PAGE evaluation (Calvete, 2011, 2014). Peptide ratio were semi-quantified by their MS-TIC signal. The UV_214nm_ peak integrals were normalized the total of peak integrals. Possible co-eluents were regarded to the stained bands density ratio based on the SDS-PAGE fractions lanes.

### 2.10 Data accessibility

Mass spectrometry proteomics data (.mgf, .raw and output files) have been deposited at the ProteomeXchange Consortium (Vizcaíno et al., 2014) (http://proteomecentral.proteomexchange.org) via the MassIVE partner repository under the project name “Venomics of the horned desert viper (*Cerastes cerastes*) and the mangrove pit viper (*Cryptelytrops purpureomaculatus*)” and data set identifier: PXD010706.

## 3 RESULTS

In this study, crude venom and single venom fractions of two vipers from different locations were studied in terms of *in vitro* cytotoxicity. To provide further insights into the active fractions of these species, we investigated the venom proteomes by using bottom-up and intact mass profiling.

### 3.1 Cytotoxicity

Cytotoxicity assays were performed on various human cancerous (SHSY5Y, MCF-7, HeLa, A-549, CaCo-2, MDA-MB-231, 253-JBV, U87MG) and one non-cancerous cell line (HEK-293) with crude venoms of *C. cerastes* and C. purpureomaculatus (see **Figure 2**).

**Figure 2:**
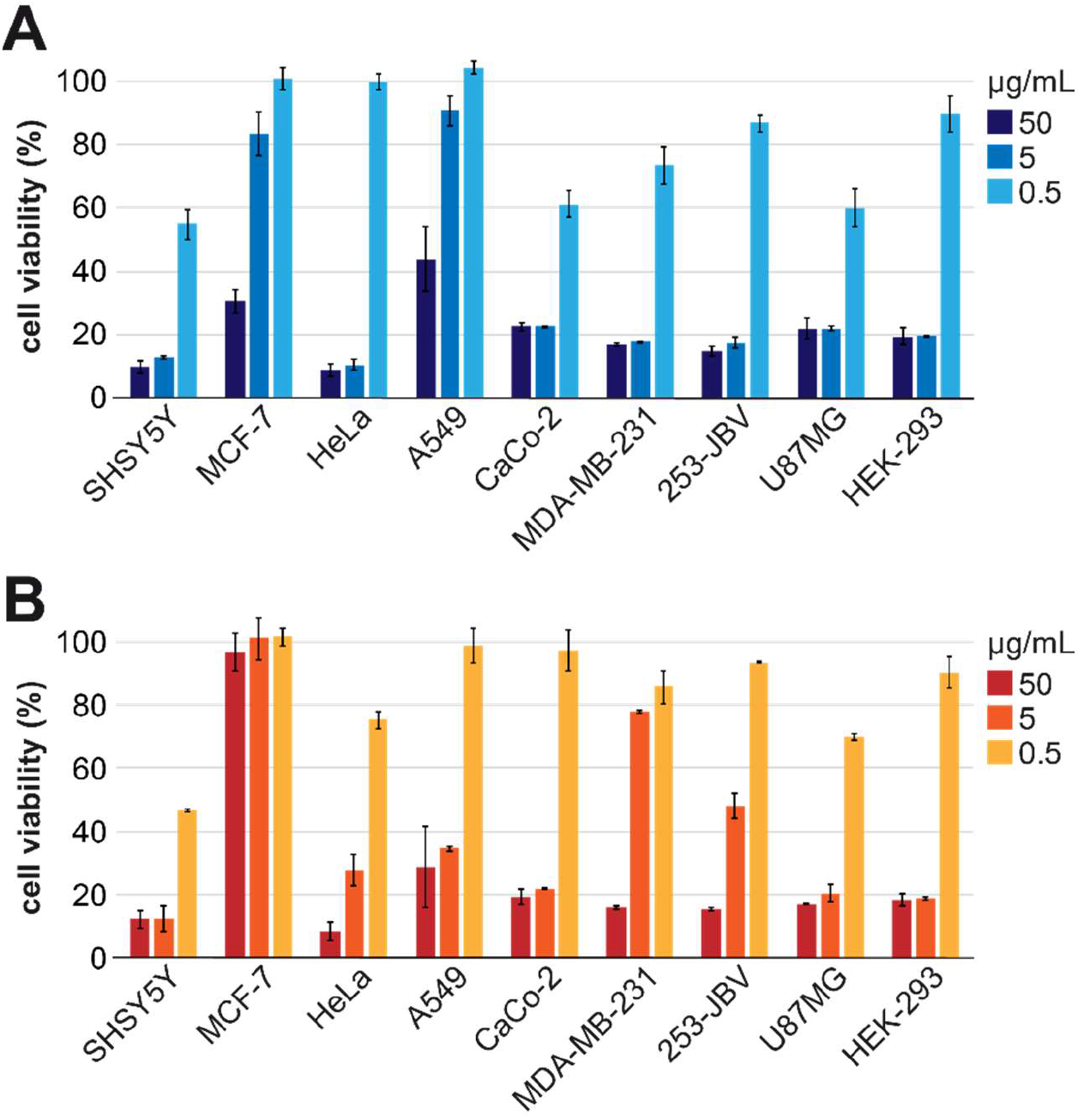
Cell viability against crude snake venoms of *Cerastes cerastes* and *Cryptelytrops purpureomaculatus*. Viability of eight cancerous human cell lines (SHSY5Y, MCF-7, HeLa, A549, CaCo-2, MDA-MB-231, 253-JBV, U87MG) and a non- cancerous cell line (HEK293) after treatment with (A) *C. cerastes* and (B) *C. purpureomaculatus* crude venom for 48 h. Cell viability was determined in an MTT assay, normalized to an untreated control (100% viability).

The 48h venom treatment shows for both vipers the most potent activity on the SHSY5Y cell line, with IC_5_ values of 0.12µg/mL and 0.25µg/mL, respectively (see **Table 1**). The venom of *C. cerastes* has higher activity levels on HeLa, CaCo-2, 253-JBV, U87MG, MDA-MB-231 and HEK-293 cell lines, whereas it showed only a moderate level of potency on A-549 and MCF-7 cell lines (see **Table 1**). However, the crude venom of *C. purpureomaculatus* showed considerable potential activity on U87MG, MDA-MB-231, HeLa, CaCo-2, A-549 and HEK-293 cell lines, with no relevant potency on the MCF-7 cell line with >50 µg/mL (see **Table 1**). In comparison, the concentration-dependent cytotoxicity shows lower IC50 values for *C. cerastes* in contrast to *C. purpureomaculatus* (see **Table 2**).

**Table 1.**
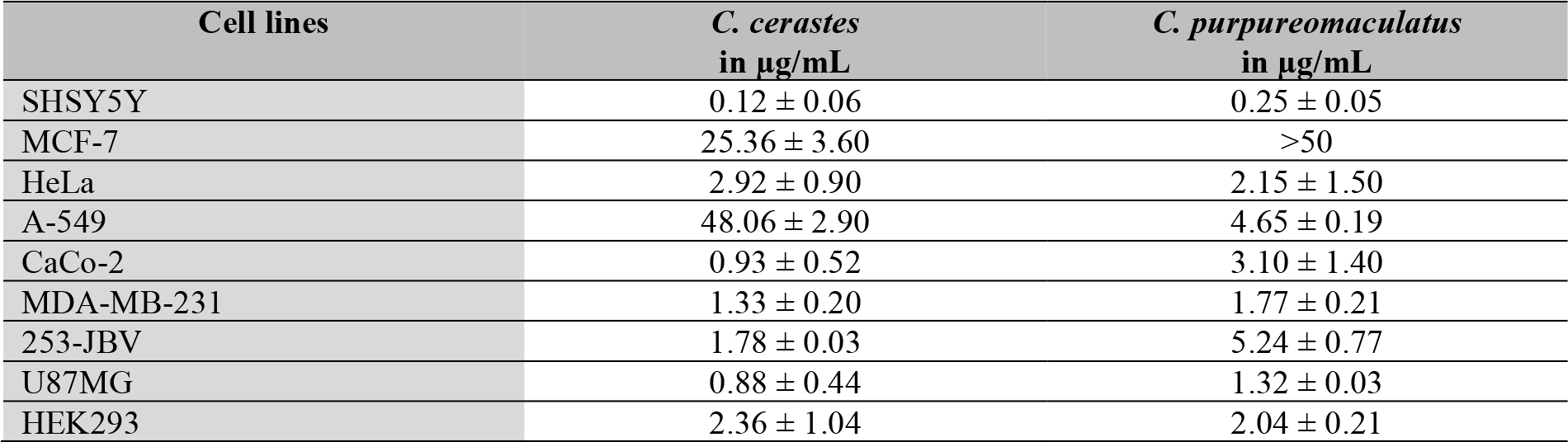
IC_50_ values of crude venoms on eight cancerous cell lines (SHSY5Y, MCF-7, HeLa, A549, CaCo-2, MDA-MB-231, 253-JBV, U87MG) and a normal cell line (HEK293) after treatment for 48 h.

Further, in-depth studies were performed with single purified venom fractions obtained from the crude venoms. The bioactivity of *C. cerastes* and *C. purpureomaculatus* venom fractions were screened against the SHSY5Y cell line, which shows the highest activity for both in the initial cytotoxicity screening (see **Figure 3**). Among the tested fractions (F) of *C. cerastes*, F8 (disintegrin), F9 (disintegrin) and F10 (predicted disintegrin) were detected as most active against SHSY5Y cells with IC_50_ values of 0.6 µM, 0.4µM and 0.1 µM, respectively (see **Figure 3**, **Table 2**). The cytotoxicity assays correlate with the morphological study as F10 had the lowest IC_50_ value with dislodged cells upon treatment (see **Figure 4**).

**Table 2.**
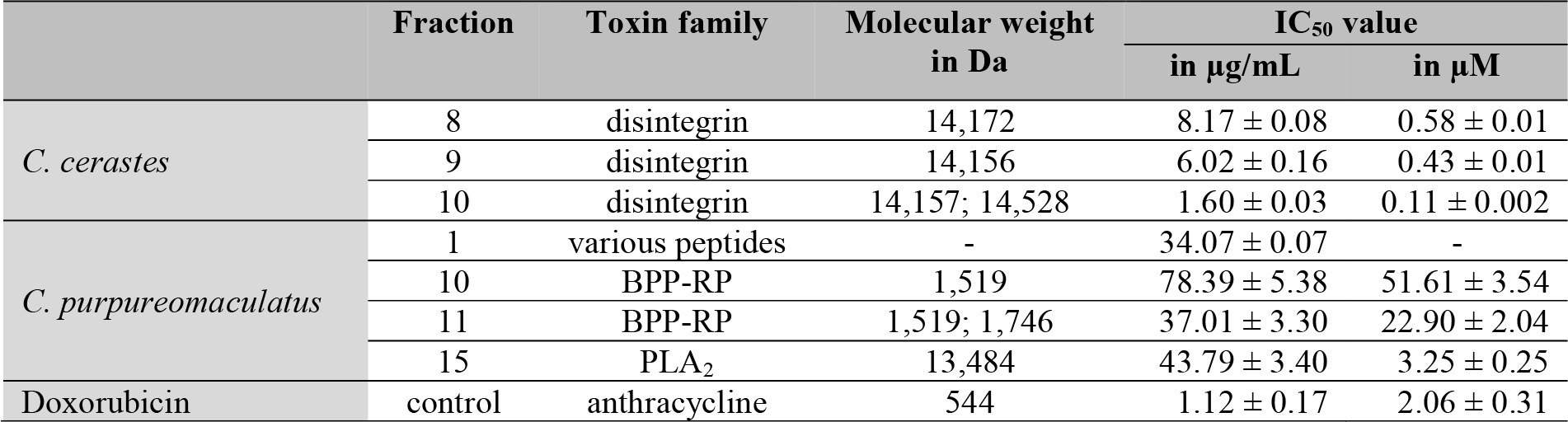
Identification of most active isolated venom fractions and IC_50_ values on SHSY5Y cell line after treatment for 48 h. In the case of two main masses in one fraction, the TIC ratio was considered for µM calculation.

**Figure 3:**
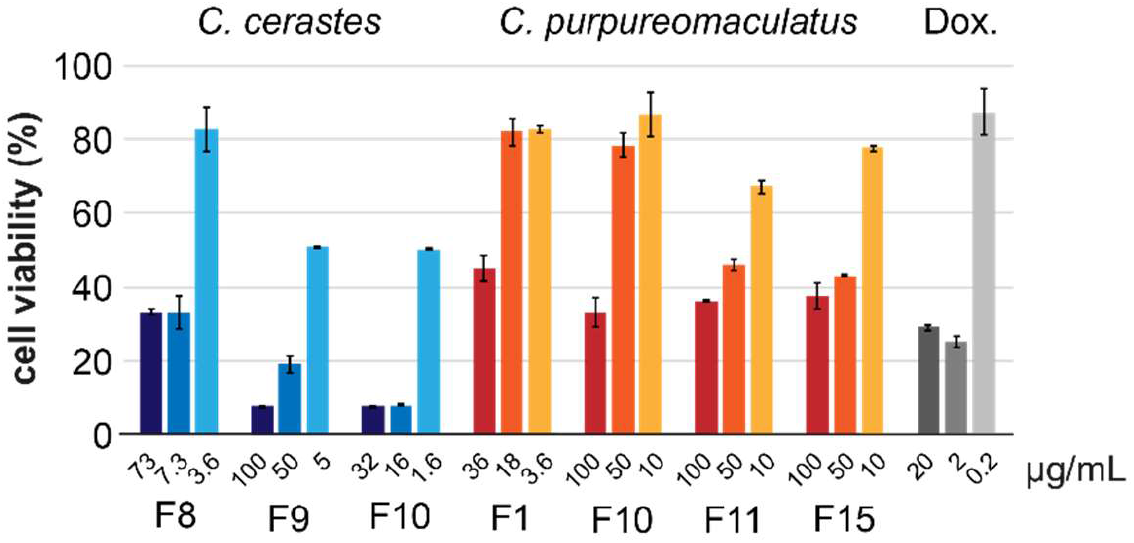
Single venom fraction cytotoxicity. Viability graphs of SHSY5Y cells after treatment with three concentrations of *C. cerastes* venom fractions (blue; F8 disintegrin, F9 disintegrin, F10 predicted disintegrin) and *C. purpureomaculatus* venom fractions (red; F1 peptides, F10 1.5 kDa peptide, F11 1.7 kDa peptide, F15 PLA2) as well as Doxorubicin as positive control (grey, Dox.) after 48 h treatment. Cell viability was determined by MTT assay and normalized to untreated control (100% viability).

**Figure 4:**
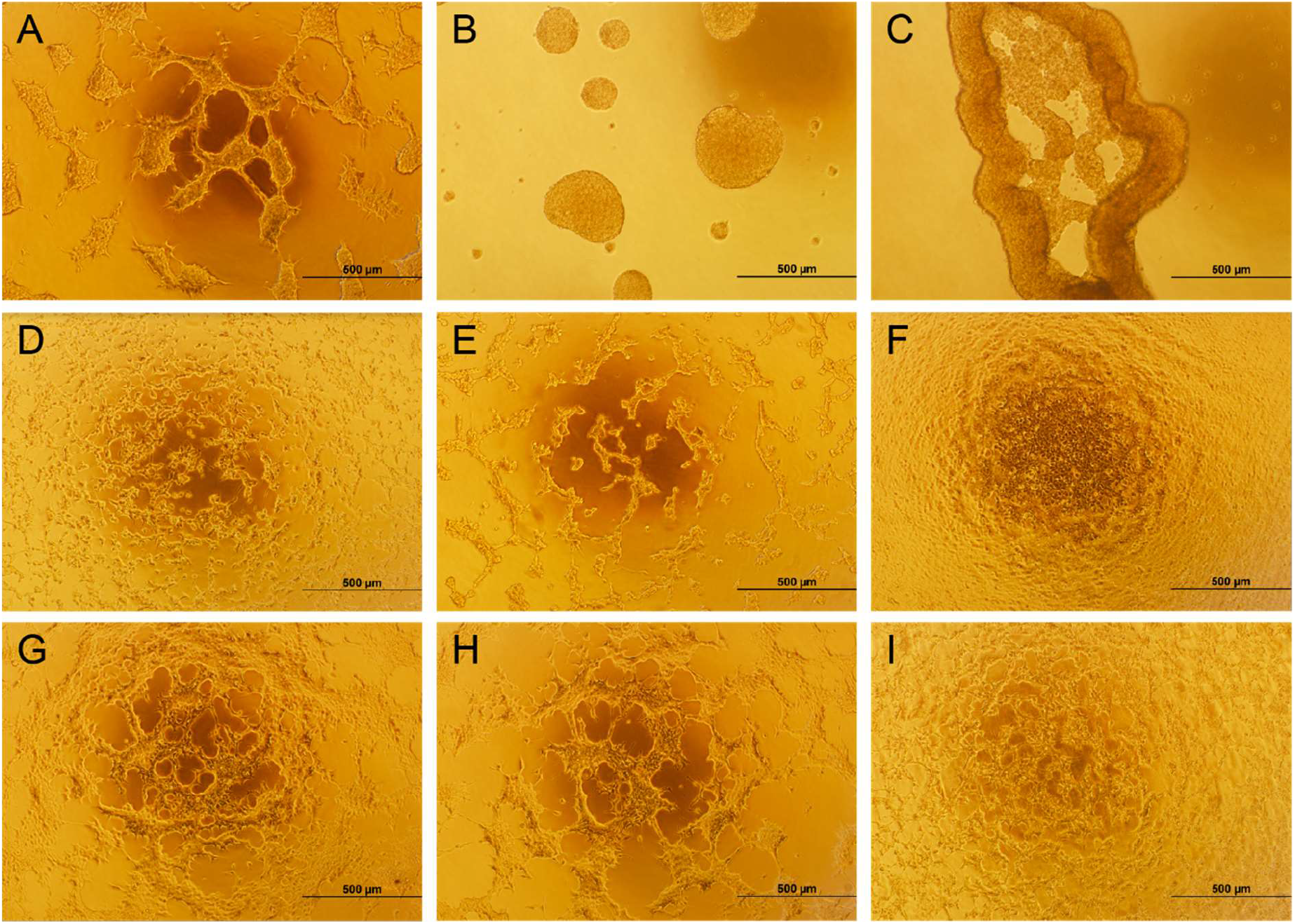
Single venom fraction of *Cerastes cerastes* and *Cryptelytrops purpureomaculatus* against SHSY5Y cells. Effect of snake venom against SHSY5Y cells upon exposure for 48 h at 37 °C. Fractions from *C. cerastes* were tested: (A) F8 (disintegrin; 7.3 µg/mL, 0.52 µM), (B) F9 (disintegrin; 5 µg/mL, 0.35 µM), (C) F10 (predicted disintegrin; 16 µg/mL, 1.13 µM). Fractions from *C. purpureomaculatus* were tested: (D) F10 (1.5 kDa peptide; 100 µg/mL, 65.83 µM), (E) F11 (1.5 and 1.7 kDa peptide; 100µg/mL, 61.87 µM), and (F) F15 (PLA2; 100 µg/mL, 7.42 µM). Doxorubicin were used as positive cytotoxic control with (G) 2 µg/mL (3.68 µM) and (H) 20 µg/mL (36.76 µM) as well as (I) an untreated control.

*C. purpureomaculatus* venom fractions demonstrated lower cytotoxicity when compared to *C. cerastes* venom fractions on the tested cell line (see **Figure 4**). In comparison, among the tested fractions of *C. purpureomaculatus*, F1 (peptide), F10 (∼1.5 kDa peptide), F11 (∼1.7 kDa peptide) and F15 (PLA_2_) were the most active fractions with cytotoxic effects against SHSY5Y cells with IC_50_ values of 52 µM (F10), 22 µM (F11) and 3.2 µM (F15) (see **Table 2**).

### 3.2 Venom proteome

The venom proteomes of both vipers were analyzed by a tryptic bottom-up approach with previous two-dimensional fractionation through HPLC and SDS as well as an IMP, and revealed distinctive differences in their protein and peptide composition (see **Figure 5, SI-Figure S1, S2**). The detailed annotation and molecular masses of venom components are shown (see **SI-Table S1, S2**). The Egypt *C. cerastes* venom composition is dominated by ∼25% of thrombin-like enzymes from the svSP family and a diversity of svMPs (∼23%) (see **Figure 5A**). Furthermore, only three other protein families were identified: 16% PLA_2_, 9% CTL, and 8% DI. The DIs could be identified in fraction F8 (0.9%) and F9 (6.9%), which includes the prevalent average mass of 14,156 Da and comparable to the theoretical average mass of a known *C. cerastes* disintegrin heterodimer CC8A+CC8B (NCBI accession no. **P83043, P83044**) of 14,154 Da with reduced Cys (Calvete et al., 2002). PLA_2_s were only identified in a single peak (F12) and form with an average mass of 13,726 Da nearly 16% of the whole venom. The rate of non-annotated proteins (10%) is relatively high, but is mainly based on F10 (∼2%) and F27 (∼7%), that includes no detectable proteins in the SDS-PAGE (see **SI-Figure S1**). In comparison to closely related fractions and the IMP, it can be assumed that F10 (14,528 Da) is another disintegrin dimer and F27 (24,104 Da) is formed by svMPs, respectively to F9 and F26.

**Figure 5:**
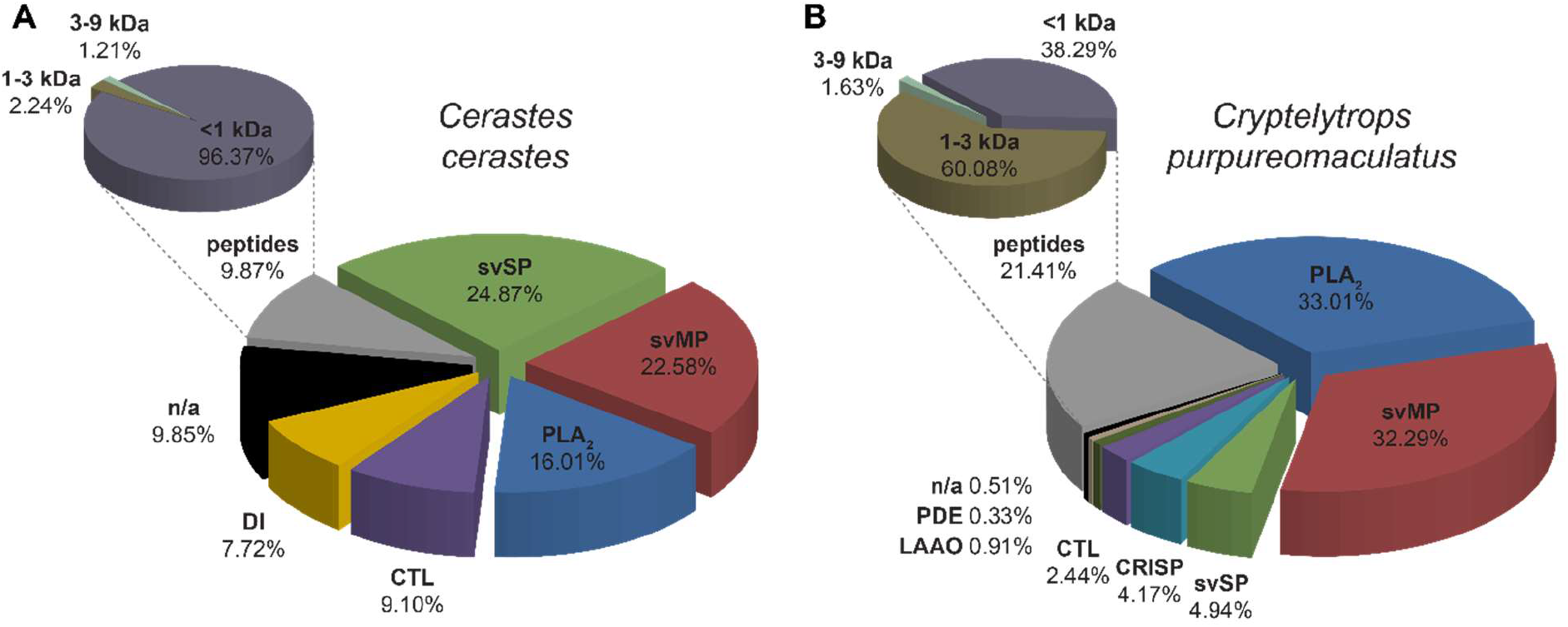
Semi-quantitative venom composition of *Cerastes cerastes* (Egypt) and *Cryptelytrops purpureomaculatus* (Thailand). The relative occurrence of different toxin families and peptide content of (A) *C. cerastes* and (B) *C. purpureomaculatus* are represented by snake venomics (λ=214 nm). We identified snake venom serine proteases (svSP, light green), snake venom metalloproteinases (svMP, red), phospholipases A_2_ (PLA_2_, blue), C-type lectin-like proteins (CTL, violet), disintegrins (DI, yellow), cysteine rich secretory proteins (CRISP, light blue), L-amino acid oxidases (LAAO, dark green), phosphodiesterase (PDE, light brown) and unknown proteins (n/a, black). Peptides (grey) grouped to different sizes are summarized in an additional pie chart percental related to the total peptide content. They were clustered to <1 kDa (dull purple), 1-3 kDa (dull brown) and 3-9 kDa (dull green) parts.

The second analyzed venom originates from a Thai *C. purpureomaculatus* and the venomic composition is shown (see **Figure 5B**). The most abundant protein family is represented by 33% PLA2, with the two main toxins in F15 (17%) and F16 (9%). The masses were identified by IMP with 13,484 Da and 13,898 Da, respectively. Next, 32% svMP is followed as the second most abundant family and svSP (5%), cysteine rich secretory proteins (CRISP, 4%) as well as CTL (2%) as minor groups. The only CRISP containing fraction (F22), was identified through a high fragment coverage against the N-terminus of the *C. purpureomaculatus* CRISP tripurin (NCBI accession no. **P81995**), which sequence is only described by 33 amino acids (Zainal Abidin et al., 2016). This protein could be identified as the main venom CRISP of *C. purpureomaculatus* with a molecular weight of 25,179 Da. The lowest abundant proteins are formed by L-amino acid oxidases (LAAO, 0.9%), phosphodiesterase (PDE, 0.3%) and the part of unidentified proteins (0.5%). In contrast to *C. cerastes* no disintegrins could be identified in the venom of *C. purpureomaculatus* by the BU approach, only in fraction F11 a molecular mass of *m/z* 7,561 was observed, which might be an indicator of low concentrations of DI.

In total, seven isolated venom fractions of both vipers were identified as most cytotoxic against SHSY5Y cells, with remarkable IC_50_ differences between the species (see **Table 2**). In combination with the BU approach, the IMP shows that the active *C. cerastes* venom fractions F8-10 includes four specimens of dimeric DI as mentioned before like the CC8A+CC8B heterodimer. All masses are in the range of 14 kDa (see **Figure 8A-C**).

The venom fractions of *C. purpureomaculatus* are less active and more divers in the corresponding families. The four fractions F1, F10, F11 and F15 are formed by several peptides and one PLA_2_ (see **Table 2**). The fractions F10 and F11 contain as main part related peptides of bradykinin-potentiating peptides (BPP-RP), which were *de novo* identified by MS2 measurements as the 13mer pEPPHWPPPHHIPP and the 16mer PPPPPPPWSPPHHIPP (see **Figure 7F-G**). Both peptides have a high sequence homology to the ‘bradykinin-potentiating and C-type natriuretic peptides’ (NCBI accession no. **P0C7P6**) of the bamboo pit viper (Trimeresurus gramineus), a related species to *C. purpureomaculatus*, former also *Trimeresurus sp.* (Malhotra and Thorpe, 2004). The BPP-RP 13mer is present in both fractions, while the 16mer has been identified from F11. As previously observed in our studies (Damm et al., 2018; Göçmen et al., 2015a; Hempel et al., 2018) Fraction F1 is composed of a multitude of small peptides and F15 contains the main PLA_2_ (13,484 Da) of the venom (see **Figure 7D-E**).

## 4 Discussion

Cytotoxic activities of snake venoms, show variations between species and tested cell lines, which could be linked in previous studies to variable modes of actions and receptor interactions of the molecules, making them attractive sources of interest for cancer therapy (Bradshaw et al., 2016; Calderon et al., 2014; Ozen et al., 2015; Yalcın et al., 2014). In the present study, crude venom of *C. cerastes* from North Africa and *C. purpureomaculatus* from Asia were screened for cytotoxicity by performing a MTT assay on eight different cancerous cell lines together with non-cancerous cell lines.

Compared to our results, previous studies on closer related members of the *Viperidae* family (*Montivipera bulgardaghica, Montivipera raddei* and *Protobothrops flavoviridis*) indicates lower cytotoxicity against comparable cell lines (Damm et al., 2018; Nalbantsoy et al., 2017). Furthermore, the venom of the related Indian Russel’s viper (*Vipera russelli*) was tested for anticancer activity in an animal model of carcinoma, and reported increased lifespan of mice, as well as cytotoxicity on leukemic cells *in vitro* (Debnath et al., 2007). A study on *C. cerastes* venom treated MCF-7 cells demonstrated a lower IC_50_ value (1.5 µg/mL) compared to this study (25.4 μg/mL) (Shebl et al., 2012). Additionally, the anti-cancer activity of an isolated LAAO from the close related *Cerastes vipera* venom against MCF-7 cells was reported with an IC_50_ value of 2.8 μg/mL (Salama et al., 2018).

In a study by Zainal Abidin et al. (2018) the cytotoxic, anti-proliferative and apoptotic activity of a LAAO from the Malaysian *C. purpureomaculatus* was screened against colon cancer cells (SW480 and SW620). They reported an EC_50_ value of 13 µg/mL against colon cancer cell lines and demonstrated morphological changes as well as apoptosis (Zainal Abidin et al., 2018). In our study, a direct comparison of the concentration-dependent cytotoxicity shows that IC_50_ values of *C. cerastes* were lower in contrast to *C. purpureomaculatus* (see **Table 2**).

In-depth cytotoxicity studies on SHSY5Y cells with purified venom fractions of *C. cerastes* and *C. purpureomaculatus* were studied here for the first time. The results obtained from disintegrin-containing fractions, accord to the disorganization and detachment of SHSY5Y cells and aggregate formation after *C. cerastes* venom fractions treatment, compared to the untreated samples, which are homogenously adhered (see **Figure 4**). Figure 4 demonstrate the typical activity of disintegrin-like proteins by dislodged and detached forming sphere morphologies as disintegrins are known for their activities on integrins, critical cell surface receptors for cell adhesion, migration, and growth, preventing cancer metastasis and growth (see *Figure 4A-C*) (Koh and Kini, 2012; Selistre-de-Araujo et al., 2010). SHSY5Y is known for its high metastatic potential achieved by β1-integrin and the effect of snake venom disintegrins could be responsible from the morphological change mentioned in Figure 4 (Meyer et al., 2004). Based on the morphological analysis, F10 as disintegrin-like peptide was found to be most potent component against SHSY5Y cells (see **Figure 4**)

The fractions of *C. purpureomaculatus* demonstrated lower cytotoxicity when compared to *C. cerastes.* The PLA_2_ activities of *C. purpureomaculatus* correlate with a previous study, which are important in membrane phospholipid hydrolysis (Chu et al., 2009; Zainal Abidin et al., 2016). These PLA_2_s mainly release fatty acids, like arachidonic acid, and lysophosholipids of the plasma membrane that are important auto- and paracrine mediators during cancer progression, although PLA_2_s from *C. purpureomaculatus* in this study showed moderate cytotoxicity (IC_50_=3.3 μM).(Ma et al., 2017). The levels of potency in *C. purpureomaculatus* venom are found to be moderate compared to the *C. cerastes* venom (see **Table 2**). These results demonstrated interesting outcomes with regard to pharmacological aspects as well as future studies on anticancer agents. In contrast to our proteomic study of the Egypt specimen, previous snake venomic analyzes about specimens of two further *C. cerastes* habitats (Morocco and Tunisia), revealed in Morocco svMP with two-third as main and PLA_2_s as second abundant family (19%), while svSPs were a minor group (7%) (Fahmi et al., 2012) (see **Figure 6**). Two Tunisian *C. cerastes* studies corroborate the diverse venom composition of this widely spread species (Bazaa et al., 2005; Fahmi et al., 2012). Both showed svMPs as most abundant family (37-56%) and were strongly divergent in their CTL content (2-24%). In analogy are the svSPs with 9-13% only less represented, compared to 25% in the venom proteome of here presented Egyptian *C. cerastes.* Therefore, the Moroccan specimen remains the only population with observed venomic CRISP proteins (Fahmi et al., 2012). This underlines the variety of the venom compositions within a single species, which is often be predicted by subpopulations and environmental influences (Amazonas et al., 2018; Daltry et al., 1996). These local differences might be important for pharmacological aspects and should be considered for the design of antivenoms. Especially since the distribution area of *C. cerastes* is wide larger than the three until now tested habitats, which reaches from Mauritania up to Iraq and Oman (GBIF Backbone Taxonomy, 2017).

**Figure 6:**
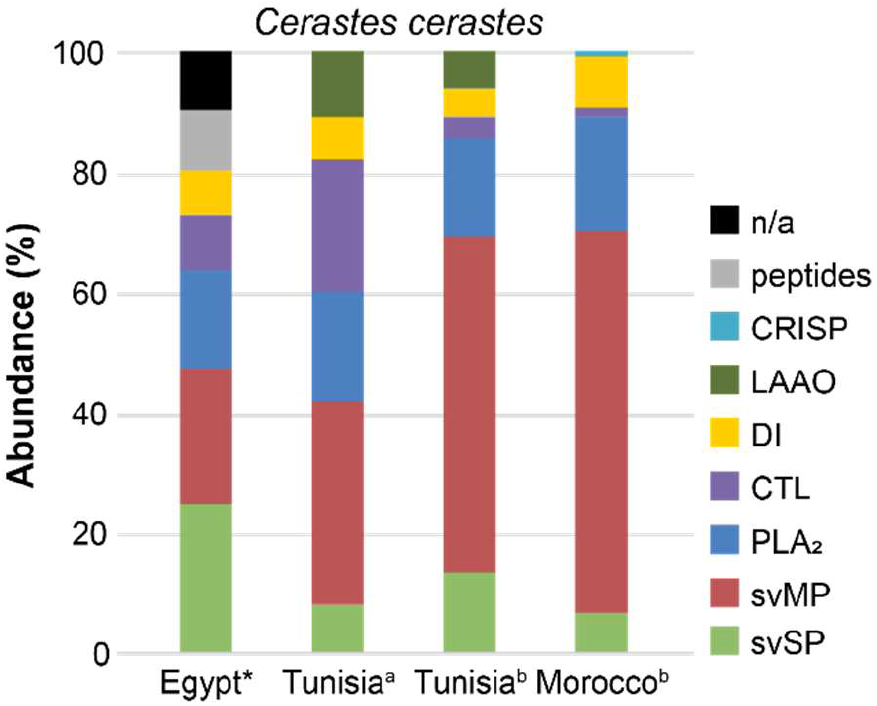
Venomic data of *Cerastes cerastes* comparison different habitats. Venom compositions of proteomic venom *C. cerastes* analysis (quantified at λ=214 and 215 nm) of three different North African natural habitats. The asterisked data set (Egypt) is the result of this study. The origins of toxin ratios are marked alphabetically: a (Bazaa et al., 2005), b (Fahmi et al., 2012).

**Figure 7:**
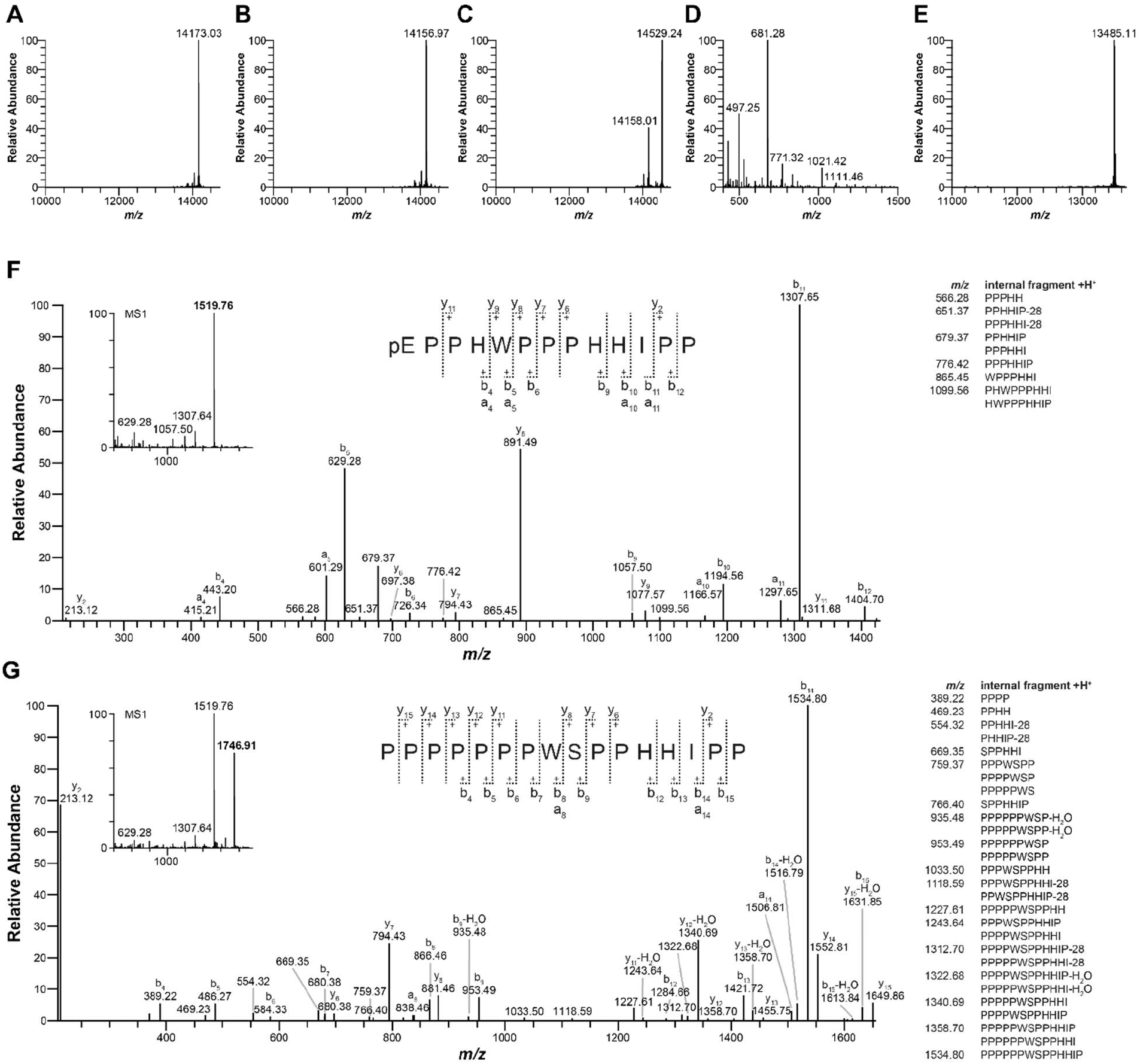
MS identification of the most active venom fractions from *Cerastes cerastes* and *Cryptelytrops purpureomaculatus*. The highest cytotoxicity against SHSY5Y cell were shown by several venom fractions with distinct molecular masses measured at the MS1 level of dimeric disintegrins for *C. cerastes* isolated fractions (A) F8 - *m/z* 14173, (B) F9 - *m/z* 14157 and (C) F10 - *m/z* 14158/14529 and for *C. purpureomaculatus* (D) F1 - *m/z* 681 and (E) F15 - *m/z* 13485. Additionally, for *C. purpureomaculatus* two BPP related peptides in (F) F10 and (G) F11 were identified and annotated at the MS2 level as pEPPHWPPPHHIPP (m/z 1520, pE as pyroglutamate) and PPPPPPPWSPPHHIPP (m/z 1747).

In the case of the *C. purpureomaculatus* a qualitative shotgun proteomic composition is known from a Malaysian species. While a direct quantitative comparison based on different analysis methods is not recommended, at the qualitative level it shows the same identified protein families as here described (Zainal Abidin et al., 2016). The lack of phospholipase B and glutaminyl-peptide cyclotransferase in this study may be an effect of the low abundance of these protein families or simply the fact that the Thai *C. purpureomaculatus* did not secrete these toxins. Furthermore, the *C. purpureomaculatus* venom includes the two mentioned dominant BPP related peptides pEPPHWPPPHHIPP and PPPPPPPWSPPHHIPP, which amino acid sequences are homologous to the general BPP structure pattern of pE-X_n_-P-X_n_-P-X_n_-IPP (with X any amino acid, except Cys) (Sciani and Pimenta, 2017). The only exception is the missing pyroglutamate at the N-terminus of the 16mer, formed by a polyproline-motif of seven prolines. Beside the internal variation of two amino acids, could this extensive terminal difference be a critical point in the changed fraction activity and its effect should be clarified by further studies.

## 5 Conclusion

In summary, two species of the *Viperidae* family, *C. cerastes* and *C. purpureomaculatus* were screened for their cytotoxicity potential against eight cancerous and one non-cancerous cell line. The cytotoxicity potentials of the venoms were effective on the tested cell lines except MCF-7 and both were found to be highly effective against SHSY5Y cell line. *C. cerastes* venom is rich in active disintegrin fractions and exerts a higher cytotoxic effect on all cell lines compared to the *C. purpureomaculatus*. Additionally, in further studies we tested venom fractions for their cytotoxicity activity. For *C. cerastes* three disintegrin fractions and for *C. purpureomaculatus* a peptide fraction, two BPP-RP (pEPPHWPPPHHIPP, PPPPPPPWSPPHHIPP) and a PLA_2_ was detected as high active fractions.

A variety of cytotoxicity studies demonstrates that snake venoms are a promising natural source with an enormous potential in terms of cancer treatment. However, our results show obstacles for being used as a therapeutic. Some of these are the toxicity against healthy cells like the HEK-293, defining the exact administration dose as well as the limited access (Vyas et al., 2013). These problems could be overcome with alternative approaches for example targeted delivery systems, identification and modification of active components from venoms and heterologous production of active components. Furthermore, we analyzed and quantified both venoms by comprehensive snake venomics. This study provides data for future studies in assessing the potential of viper venoms in anti-cancer studies as the highly active fractions were found especially in the *C. cerastes* venom.

## Acknowledgement

We would like to thank Joshua Baal for providing us with images of the Egyptian horned desert viper (*Cerastes cerastes*) and Thai mangrove pit viper (*Cryptelytrops purpureomaculatus*). Additionally, we would like to thank Mert Karıs and Mehmet Anıl Oguz of the Zoology Section from Ege University (Izmir, Turkey) for participating the venom extraction process.

## Authors contribution

B.-F.H., and A.N. planned the study. B.G., M.K. and M.A.O. extracted the crude snake venoms. B.-F.H., and R.S. performed the protein separation and acquired the mass spectrometry data. A.N. and C.S.O performed the cytotoxicity screenings. B.-F.H., M.D., C.S.O and R.S. performed the data analysis. A.N. and R.D.S. acquired funding and provided materials and instruments for the study. B.-F.H., M.D., C.S.O and R.D.S. wrote the manuscript. All authors read, discussed and approved the manuscript.

## References

Aird, S.D., 2005. Taxonomic distribution and quantitative analysis of free purine and pyrimidine nucleosides in snake venoms. Comparative biochemistry and physiology. Part B, Biochemistry & molecular biology 140 (1), 109–126.

Al-Sadoon, M.K., Paray, B.A., 2016. Ecological aspects of the horned viper, *Cerastes cerastes gasperettii* in the central region of Saudi Arabia. Saudi journal of biological sciences 23 (1), 135–138.

Amazonas, D.R., Portes-Junior, J.A., Nishiyama-Jr, M.Y., Nicolau, C.A., Chalkidis, H.M., Mourão, R.H.V., Grazziotin, F.G., Rokyta, D.R., Gibbs, H.L., Valente, R.H., Junqueira-de-Azevedo, I.L.M., Moura-da-Silva, A.M., 2018. Molecular mechanisms underlying intraspecific variation in snake venom. Journal of proteomics 181, 60–72.

Antunes, T.C., Yamashita, K.M., Barbaro, K.C., Saiki, M., Santoro, M.L., 2010. Comparative analysis of newborn and adult *Bothrops jararaca snake venoms*. Toxicon: official journal of the International Society on Toxinology 56 (8), 1443–1458.

Barlow, A., Pook, C.E., Harrison, R.A., Wüster, W., 2009. Coevolution of diet and prey-specific venom activity supports the role of selection in snake venom evolution. Proceedings. Biological sciences 276 (1666), 2443–2449.

Bazaa, A., Marrakchi, N., El Ayeb, M., Sanz, L., Calvete, J.J., 2005. Snake venomics: Comparative analysis of the venom proteomes of the Tunisian snakes *Cerastes cerastes, Cerastes vipera* and *Macrovipera lebetina*. Proteomics 5 (16), 4223–4235.

Biardi, J.E., Chien, D.C., Coss, R.G., 2006. California ground squirrel (*Spermophilus beecheyi*) defenses against rattlesnake venom digestive and hemostatic toxins. Journal of chemical ecology 32 (1), 137–154.

Bradford, M.M., 1976. A rapid and sensitive method for the quantitation of microgram quantities of protein utilizing the principle of protein-dye binding. Analytical biochemistry 72, 248–254.

Bradshaw, M.J., Saviola, A.J., Fesler, E., Mackessy, S.P., 2016. Evaluation of cytotoxic activities of snake venoms toward breast (MCF-7) and skin cancer (A-375) cell lines. Cytotechnology 68 (4), 687–700.

Calderon, L.A., Sobrinho, J.C., Zaqueo, K.D., Moura, A.A. de, Grabner, A.N., Mazzi, M.V., Marcussi, S., Nomizo, A., Fernandes, C.F.C., Zuliani, J.P., Carvalho, B.M.A., da Silva, S.L., Stábeli, R.G., Soares, A.M., 2014. Antitumoral activity of snake venom proteins: New trends in cancer therapy. BioMed research international 2014, 203639.

Calvete, J.J., 2011. Proteomic tools against the neglected pathology of snake bite envenoming. Expert review of proteomics 8 (6), 739–758.

Calvete, J.J., 2014. Next-generation snake venomics: Protein-locus resolution through venom proteome decomplexation. Expert review of proteomics 11 (3), 315–329.

Calvete, J.J., Fox, J.W., Agelan, A., Niewiarowski, S., Marcinkiewicz, C., 2002. The presence of the WGD motif in CC8 heterodimeric disintegrin increases its inhibitory effect on alphaII(b)beta3, alpha(v)beta3, and alpha5beta1 integrins. Biochemistry 41 (6), 2014–2021.

Casewell, N.R., Wüster, W., Vonk, F.J., Harrison, R.A., Fry, B.G., 2013. Complex cocktails: The evolutionary novelty of venoms. Trends in ecology & evolution 28 (4), 219–229.

Chan, Y.S., Cheung, R.C.F., Xia, L., Wong, J.H., Ng, T.B., Chan, W.Y., 2016. Snake venom toxins: Toxicity and medicinal applications. Applied microbiology and biotechnology 100 (14), 6165–6181.

Chijiwa, T., Deshimaru, M., Nobuhisa, I., Nakai, M., Ogawa, T., Oda, N., Nakashima, K., Fukumaki, Y., Shimohigashi, Y., Hattori, S., Ohno, M., 2000. Regional evolution of venom-gland phospholipase A2 isoenzymes of *Trimeresurus flavoviridis* snakes in the southwestern islands of Japan. The Biochemical journal 347 (Pt 2), 491–499.

Chippaux, J.P., Williams, V., White, J., 1991. Snake venom variability: Methods of study, results and interpretation. Toxicon: official journal of the International Society on Toxinology 29 (11), 1279–1303.

Chu, C.-W., Tsai, T.-S., Tsai, I.-H., Lin, Y.-S., Tu, M.-C., 2009. Prey envenomation does not improve digestive performance in Taiwanese pit vipers (*Trimeresurus gracilis* and *T. stejnegeri stejnegeri*). Comparative biochemistry and physiology. Part A, Molecular & integrative physiology 152 (4), 579–585.

Daltry, J.C., Wüster, W., Thorpe, R.S., 1996. Diet and snake venom evolution. Nature 379 (6565), 537–540.

Damm, M., Hempel, B.-F., Nalbantsoy, A., Süssmuth, R.D., 2018. Comprehensive Snake Venomics of the Okinawa Habu Pit Viper, *Protobothrops flavoviridis*, by Complementary Mass Spectrometry-Guided Approaches. Molecules (Basel, Switzerland) 23 (8).

Das, T., Bhattacharya, S., Biswas, A., Gupta, S.D., Gomes, A., Gomes, A., 2013. Inhibition of leukemic U937 cell growth by induction of apoptosis, cell cycle arrest and suppression of VEGF, MMP-2 and MMP-9 activities by cytotoxin protein NN-32 purified from Indian spectacled cobra (Naja naja) venom. Toxicon: official journal of the International Society on Toxinology 65, 1–4.

Debnath, A., Chatterjee, U., Das, M., Vedasiromoni, J.R., Gomes, A., 2007. Venom of Indian monocellate cobra and Russell’s viper show anticancer activity in experimental models. Journal of ethnopharmacology 111 (3), 681–684.

Dekhil, H., Wisner, A., Marrakchi, N., El Ayeb, M., Bon, C., Karoui, H., 2003. Molecular cloning and expression of a functional snake venom serine proteinase, with platelet aggregating activity, from the Cerastes cerastes viper. Biochemistry 42 (36), 10609–10618.

Devi, A., 2013. The Protein and Nonprotein Constituents of Snake Venoms. In: Bücherl, W., Buckley, E.E., Deulofeu, V. (Eds.), Venomous Animals and Their Venoms. Venomous Vertebrates. Elsevier Science, Burlington, pp. 119–165.

Fahmi, L., Makran, B., Pla, D., Sanz, L., Oukkache, N., Lkhider, M., Harrison, R.A., Ghalim, N., Calvete, J.J., 2012. Venomics and antivenomics profiles of North African *Cerastes cerastes* and *C. vipera* populations reveals a potentially important therapeutic weakness. Journal of proteomics 75 (8), 2442–2453.

Fry, B.G., 2015. Venomous reptiles and their toxins: Evolution, pathophysiology, and biodiscovery. Oxford University Press, New York, NY.

GBIF Backbone Taxonomy, 2017. *Cerastes cerastes* Linnaeus, 1758 in GBIF Secretariat (2017): Checklist dataset. Accessed 16 October 2018.

Gibbs, H.L., Mackessy, S.P., 2009. Functional basis of a molecular adaptation: Prey-specific toxic effects of venom from *Sistrurus* rattlesnakes. Toxicon: official journal of the International Society on Toxinology 53 (6), 672–679.

Göçmen, B., Heiss, P., Petras, D., Nalbantsoy, A., Süssmuth, R.D., 2015a. Mass spectrometry guided venom profiling and bioactivity screening of the Anatolian Meadow Viper, *Vipera anatolica*. Toxicon: official journal of the International Society on Toxinology 107 (Pt B), 163–174.

Göçmen, B., Mulder, J., Karış, M., Mebert, K., 2015b. New locality records of *Vipera ammodytes transcaucasiana* (Boulenger, 1913) in Turkey. South Western Journal of Horticulture, Biology and Eviroment 6 (2), 91–98.

Gopalakrishnakone, P., Faiz, A., Fernando, R., Gnanathasan, C.A., Habib, A.G., Yang, C.-C. (Eds.), 2015. Clinical Toxinology in Asia Pacific and Africa, vol. 2. Toxinology. Springer Netherlands, Dordrecht, s.l., 158 pp.

Greene, H.W., 1997. Snakes: The evolution of mystery in nature. Univ. of California Press, Oxford, 351 pp.

Harvey, A.L., 2014. Toxins and drug discovery. Toxicon: official journal of the International Society on Toxinology 92, 193–200.

Hempel, B.-F., Damm, M., Göçmen, B., Karis, M., Oguz, M.A., Nalbantsoy, A., Süssmuth, R.D., 2018. Comparative Venomics of the *Vipera ammodytes transcaucasiana* and *Vipera ammodytes montandoni* from Turkey Provides Insights into Kinship. Toxins 10 (1).

Jain, D., Kumar, S., 2012. Snake Venom: A Potent Anticancer Agent. Asian Pacific Journal of Cancer Prevention 13 (10), 4855–4860.

Kakanj, M., Ghazi-Khansari, M., Zare Mirakabadi, A., Daraei, B., Vatanpour, H., 2015. Cytotoxic Effect of Iranian *Vipera lebetina* Snake Venom on HUVEC Cells. Iranian journal of pharmaceutical research: IJPR 14 (Suppl), 109–114.

Kitchens, C., Eskin, T., 2008. Fatality in a case of envenomation by *Crotalus adamanteus* initially successfully treated with polyvalent ovine antivenom followed by recurrence of defibrinogenation syndrome. Journal of medical toxicology: official journal of the American College of Medical Toxicology 4 (3), 180–183.

Koh, C.Y., Kini, R.M., 2012. From snake venom toxins to therapeutics - cardiovascular examples. Toxicon: official journal of the International Society on Toxinology 59 (4), 497–506.

Konshina, A.G., Boldyrev, I.A., Utkin, Y.N., Omel’kov, A.V., Efremov, R.G., 2011. Snake cytotoxins bind to membranes via interactions with phosphatidylserine head groups of lipids. PloS one 6 (4), e19064.

Laemmli, U.K., 1970. Cleavage of structural proteins during the assembly of the head of bacteriophage T4. Nature 227 (5259), 680–685.

Leong, P.K., Tan, C.H., Sim, S.M., Fung, S.Y., Sumana, K., Sitprija, V., Tan, N.H., 2014. Cross neutralization of common Southeast Asian viperid venoms by a Thai polyvalent snake antivenom (Hemato Polyvalent Snake Antivenom). Acta tropica 132, 7–14.

Li, L., Huang, J., Lin, Y., 2018. Snake Venoms in Cancer Therapy: Past, Present and Future. Toxins 10 (9).

Ma, R., Mahadevappa, R., Kwok, H.F., 2017. Venom-based peptide therapy: Insights into anticancer mechanism. Oncotarget 8 (59), 100908–100930.

Macedo, S.R.A., Barros, N.B. de, Ferreira, A.S., Moreira-Dill, L.S., Calderon, L.A., Soares, A.M., Nicolete, R., 2015. Biodegradable microparticles containing crotamine isolated from *Crotalus durissus terrificus* display antileishmanial activity in vitro. Pharmacology 95 (1-2), 78–86.

Mackessy, S.P. (Ed.), 2010. Handbook of venoms and toxins of reptiles. CRC Press, Boca Raton, Fla., 521 pp.

Malhotra, A., Thorpe, R.S., 2004. A phylogeny of four mitochondrial gene regions suggests a revised taxonomy for Asian pitvipers (*Trimeresurus and Ovophis*). Molecular phylogenetics and evolution 32 (1), 83–100.

Marsh, N., Williams, V., 2005. Practical applications of snake venom toxins in haemostasis. Toxicon: official journal of the International Society on Toxinology 45 (8), 1171–1181.

Meyer, A., van Golen, C.M., Kim, B., van Golen, K.L., Feldman, E.L., 2004. Integrin expression regulates neuroblastoma attachment and migration. Neoplasia (New York, N.Y.) 6 (4), 332–342.

Mong, R., Tan, H.H., 2016. Snakebite by the Shore Pit Viper (*Trimeresurus purpureomaculatus*) Treated With Polyvalent Antivenom. Wilderness & environmental medicine 27 (2), 266–270.

Mosmann, T., 1983. Rapid colorimetric assay for cellular growth and survival: Application to proliferation and cytotoxicity assays. Journal of immunological methods 65 (1-2), 55–63.

Muth, T., Weilnböck, L., Rapp, E., Huber, C.G., Martens, L., Vaudel, M., Barsnes, H., 2014. DeNovoGUI: An open source graphical user interface for de novo sequencing of tandem mass spectra. Journal of proteome research 13 (2), 1143–1146.

Nalbantsoy, A., Hempel, B.-F., Petras, D., Heiss, P., Göçmen, B., Iğci, N., Yildiz, M.Z., Süssmuth, R.D., 2017. Combined venom profiling and cytotoxicity screening of the Radde’s mountain viper (*Montivipera raddei*) and Mount Bulgar Viper (*Montivipera bulgardaghica*) with potent cytotoxicity against human A549 lung carcinoma cells. Toxicon: official journal of the International Society on Toxinology 135, 71–83.

Ozen, M.O., İğci, N., Yalçin, H.T., Goçmen, B., Nalbantsoy, A., 2015. Screening of cytotoxic and antimicrobial activity potential of Anatolian *Macrovipera lebetina obtusa* (Ophidia: Viperidae) crude venom. Frontiers in Life Science 8 (4), 363–370.

Salama, W.H., Ibrahim, N.M., El Hakim, A.E., Bassuiny, R.I., Mohamed, M.M., Mousa, F.M., Ali, M.M., 2018. l-Amino acid oxidase from *Cerastes vipera* snake venom: Isolation, characterization and biological effects on bacteria and tumor cell lines. Toxicon: official journal of the International Society on Toxinology 150, 270–279.

Sales, P.B.V., Santoro, M.L., 2008. Nucleotidase and DNase activities in Brazilian snake venoms. Comparative biochemistry and physiology. Toxicology & pharmacology: CBP 147 (1), 85–95.

Schwaner, T.D., Sarre, S.D., 1988. Body Size of Tiger Snakes in Southern Australia, with Particular Reference to *Notechis ater serventyi* (Elapidae) on Chappell Island. Journal of Herpetology 22 (1), 24.

Sciani, J.M., Pimenta, D.C., 2017. The modular nature of bradykinin-potentiating peptides isolated from snake venoms. The journal of venomous animals and toxins including tropical diseases 23, 45.

Selistre-de-Araujo, H.S., Pontes, C.L.S., Montenegro, C.F., Martin, A.C.B.M., 2010. Snake venom disintegrins and cell migration. Toxins 2 (11), 2606–2621.

Shebl, R.I., Mohamed, A.F., Ali, A.E., Amin, M.A., 2012. *Cerastes cerastes* and *Vipera lebetina* Snake Venoms Apoptotic – Stimulating Activity to Human Breast Cancer Cells and Related Gene Modulation. Journal of Cancer Science & Therapy 4 (10), 317–323.

Smith, C.G., Vane, J.R., 2003. The discovery of captopril. FASEB journal: official publication of the Federation of American Societies for Experimental Biology 17 (8), 788–789.

Suzergoz, F., Iğci, N., Cavus, C., Yildiz, M.Z., Coskun, M.B., Göçmen, B., 2016. In Vitro Cytotoxic and Proapoptotic Activities of Anatolian *Macrovipera Lebetina Obtusa* (Dwigubski, 1832) Crude Venom on Cultured K562 Human Chronic Myelogenous Leukemia Cells. UHOD 26 (1).

Tan, N.H., Armugam, A., Tan, C.S., 1989. A comparative study of the enzymatic and toxic properties of venoms of the Asian lance-headed pit viper (Genus *Trimeresurus*). Comparative biochemistry and physiology. B, Comparative biochemistry 93 (4), 757–762.

Tashima, A.K., Sanz, L., Camargo, A.C.M., Serrano, S.M.T., Calvete, J.J., 2008. Snake venomics of the Brazilian pitvipers *Bothrops cotiara* and *Bothrops fonsecai*. Identification of taxonomy markers. Journal of proteomics 71 (4), 473–485.

Tomović, L., 2006. Systematics of the nose-horned viper (*Vipera ammodytes*, Linnaeus, 1758). Herpetological Journal 16 (2), 191–201.

Vanzolini, P.E., Calleffo, M.E.V., 2002. A taxonomic bibliography of the South American snakes of the *Crotalus durissus* complex (*Serpentes, Viperidae*). An. Acad. Bras. Ciênc. 74 (1), 37–83.

Vieira Santos, M.M. de, Sant’Ana, C.D., Giglio, J.R., da Silva, R.J., Sampaio, S.V., Soares, A.M., Fecchio, D., 2008. Antitumoural effect of an L-amino acid oxidase isolated from *Bothrops jararaca* snake venom. Basic & clinical pharmacology & toxicology 102 (6), 533–542.

Vizcaíno, J.A., Deutsch, E.W., Wang, R., Csordas, A., Reisinger, F., Ríos, D., Dianes, J.A., Sun, Z., Farrah, T., Bandeira, N., Binz, P.-A., Xenarios, I., Eisenacher, M., Mayer, G., Gatto, L., Campos, A., Chalkley, R.J., Kraus, H.-J., Albar, J.P., Martinez-Bartolomé, S., Apweiler, R., Omenn, G.S., Martens, L., Jones, A.R., Hermjakob, H., 2014. ProteomeXchange provides globally coordinated proteomics data submission and dissemination. Nature biotechnology 32 (3), 223–226.

Vogel, G., Grismer, L., Chan-Ard, T., 2012. Cryptelytrops purpureomaculatus. The IUCN Red List of Threatened Species 2012, e.T192188A2052944.

Vyas, V.K., Brahmbhatt, K., Bhatt, H., Parmar, U., 2013. Therapeutic potential of snake venom in cancer therapy: Current perspectives. Asian Pacific Journal of Tropical Biomedicine 3 (2), 156–162.

Yalcın, H.T., Ozen, M.O., Göçmen, B., Nalbantsoy, A., 2014. Effect of Ottoman Viper (*Montivipera xanthina* (Gray, 1849)) Venom on Various Cancer Cells and on Microorganisms. Cytotechnology 66 (1), 87–94.

Zainal Abidin, S.A., Rajadurai, P., Chowdhury, M.E.H., Ahmad Rusmili, M.R., Othman, I., Naidu, R., 2016. Proteomic Characterization and Comparison of Malaysian *Tropidolaemus wagleri* and *Cryptelytrops purpureomaculatus* Venom Using Shotgun-Proteomics. Toxins 8 (10).

Zainal Abidin, S.A., Rajadurai, P., Hoque Chowdhury, M.E., Othman, I., Naidu, R., 2018. Cytotoxic, Anti-Proliferative and Apoptosis Activity of l-Amino Acid Oxidase from Malaysian Cryptelytrops purpureomaculatus (CP-LAAO) Venom on Human Colon Cancer Cells. Molecules (Basel, Switzerland) 23 (6).

